# Metagenomic profiling pipelines improve taxonomic classification for 16S amplicon sequencing data

**DOI:** 10.1101/2022.07.27.501757

**Authors:** Tyler Faits, Aubrey R. Odom-Mabey, Eduardo Castro-Nallar, Keith A. Crandall, W. Evan Johnson

## Abstract

Most experiments studying bacterial microbiomes rely on the PCR amplification of all or part of the gene for the 16S rRNA subunit, which serves as a biomarker for identifying and quantifying the various taxa present in a microbiome sample. Several computational methods exist for analyzing 16S amplicon-based metagenomics. However, the most-used bioinformatics tools cannot produce quality genus-level or species-level taxonomic calls and may underestimate the degree to which these calls are possible. We used 16S sequencing data from mock bacterial communities to evaluate the sensitivity and specificity of several bioinformatics pipelines and genomic reference libraries used for microbiome analyses, concentrating on measuring the accuracy of species-level taxonomic assignments of 16S amplicon reads. We evaluated the tools Qiime 2, Mothur, PathoScope 2, and Kraken 2 in conjunction with reference libraries from Greengenes, Silva, Kraken, and RefSeq. Profiling tools were compared using publicly available mock community data from several sources, comprising 136 samples with varied species richness and evenness, several different amplified regions within the 16S gene, and both DNA spike-ins and cDNA from collections of plated cells. PathoScope 2 and Kraken 2, both tools designed for whole-genome metagenomics, outperformed Qiime 2 using the DADA2 plugin and Mothur, both of which are theoretically specialized for 16S analyses.

## Introduction

High-throughput sequencing has greatly accelerated the study of microbiomics. Characterizing the composition of microbial samples commonly relies on the amplification of 16S ribosomal subunit sequences, a ubiquitous gene with highly conserved regions. The subunit simplifies efforts to isolate and amplify 16S rRNA with established PCR primers and hypervariable regions to establish identity and phylogeny. 16S rRNA and rDNA sequencing can be used to identify known prokaryotic species and act as a proxy to quantify the relative abundances of operational taxonomic units (OTUs) within microbiome samples.

Methods for taxonomic profiling of ribosomal RNA gene sequences enable sample OTU identification by classifying rRNA sequences into taxonomic groups. While considerable accuracy in species-level identification is attainable with available tools^1^, current profiling software for 16S amplicon sequencing data hedges away from identifying down to the species level. Instead, they cluster reads based on sequence similarity to assign genus or higher-level identifications to increase specificity and sensitivity, or they directly use error-filtered sequences for taxonomic classification^2,3^. As the capabilities of modern sequencing platforms increase, and as bacterial reference genome databases expand and improve, more potential arises for achieving enhanced 16S analysis performance with alternative methods more commonly applied in whole genome metagenomics.

The most common software packages currently employed in analysis of 16S amplicon sequencing data are Qiime 2^4^, its predecessor, Qiime^5^, and Mothur^6^. Qiime and Mothur were both originally developed shortly after the invention of next-generation sequencing and, along with Qiime 2, essentially follow the same workflow: reads are typically clustered de novo based on sequence similarity into operational taxonomic units (OTUs) or amplicon sequence variants (ASVs) depending on whether complete sequence identity is desired for clustering. The initial clustering step serves to 1) improve computational efficiency by limiting the number of sequences needing alignment to a large set of reference genomes and 2) accommodate the low levels of genetic variation present within a given bacterial strain, thereby mitigating sequencing errors. For nearly a decade, the cutoff for OTU inclusion was 97% sequence identity^7,8^, but current cutoff recommendations are now around 99-100% sequence identity^2,9^, typically after some form of denoising or other correction for sequencing errors^3,10^.

An alternative to OTU/ASV clustering includes directly aligning reads against a reference genome library, as is done by PathoScope 2.0^11^. PathoScope employs a Bayesian mixed modeling framework to reassign ambiguously aligned reads, dampening potential sequencing errors and minor genetic variation^12,13^. As another alternative, Kraken 2 performs alignment-free *k*-mer searches against a reference genome library^14^ and makes taxonomic assignments to each read based on the cumulative number of *k*-mer matches across an entire read against each taxonomic node in its reference library. By bypassing a sequence clustering step, PathoScope and Kraken individually avoid the potential pitfalls inherent in OTU generation and denoising errors^15,16^, although remain susceptible to sequencing errors. While Qiime 2, Mothur, Greengenes, and Silva are all tools designed to address the specific needs of 16S amplicon sequencing, improvements in sequencing technologies, expanding bacterial reference genome databases, and increased availability and affordability of computational resources have collectively made many of the specific issues addressed by these tools irrelevant. Meanwhile, the increased flexibility and power of a tool such as PathoScope may yield augmented results despite being computationally intensive and designed to fulfill a more general metagenomics purpose^17,18^.

These profiling methods all rely heavily on the quality of the reference library used, as has been shown in previous benchmarking studies^19-22^. The most commonly used reference databases for 16S amplicon analyses are Greengenes^23^, Silva^24^, and the Ribosomal Database Project (RDP)^25^. Each database exclusively contains 16S gene sequences and offers taxonomic information for each reference sequence. Silva is well maintained and releases updates regularly, although as of this writing, the most recent update is Silva 138.1 (released on August 27, 2020). Meanwhile, Greengenes has been stagnant for years; its most recent update was Greengenes 13_8, released in August 2013. As a result, Greengenes lacks several essential bacteria, including *Dolosigranulum* species^26^, implicated as playing a protective role in preventing disease in human airways^27,28^. Although Qiime 2 and Mothur are compatible with any reference genome library, Qiime 2 uses Greengenes by default, and Mothur’s documentation (as accessed on May 17, 2022) recommends Silva. Kraken 2 has its own curated “Standard” bacterial library, with a taxonomic tree based on NCBI’s taxonomy database by default^29^, and has also released Kraken 2-compatible formatted versions of Greengenes, Silva, and RDP. The current PathoScope reference library recommendation is to download the complete RefSeq representative genome database^30^, a collection of curated high-quality bacterial genomes and assemblies. RefSeq is constantly updated, and as such, results of any analysis using RefSeq as a reference library may vary according to the date of the library download.

Given these considerations, we systematically benchmarked several current community profiling tools and reference libraries created for both metagenomic and 16S analysis. We evaluated the tools Qiime 2, Mothur, PathoScope 2, and Kraken 2 in conjunction with reference libraries from Greengenes, Silva, Kraken, and RefSeq. Using several publicly available 16S sequencing datasets of synthetic mock communities, we specifically analyzed genus- and species-level performance across pairs of profilers and libraries. We tested 136 samples comprising varying species richness and evenness, several different amplified regions within the 16S gene, and both DNA spike-ins and cDNA from collections of plated cells. Our evaluative comparisons utilized a combination of diversity and accuracy-based measures to determine what methods and tools provided the best performance in profiling 16S amplicon sequencing datasets.

## Methods

### Publicly available mock community sequencing datasets

136 mock community sequencing samples were collected in total from four publicly available sequencing datasets and analyzed in our evaluation. 69 samples are from Lluch et al.^31^; 33 samples are from Kozich et al.^32^; 29 samples are from Fouhy et al.^33^; and 5 samples are from Karstens^34^. These datasets are hereafter referred to as the Lluch, Kozich, Fouhy, and Karstens samples. Supplementary Table 1 delineates the species compositions for each set. The Lluch samples include a variety of community compositions, ranging from monoculture samples composed of only a single species to others with 20 species at staggered concentrations. Collectively, 34 species appear across the aggregate of Lluch samples. While the Lluch samples’ taxonomic profiles are diverse, all 69 samples were produced using a single unified DNA extraction, amplification, and sequencing protocol that yielded Illumina MiSeq paired-end reads of the V4-V5 region of the 16S gene. The Kozich samples each comprise three sequencing replicates of 11 different preparations of BEI’s mock community B (HM-278D), encompassing 21 species. For the Kozich samples, three PCR primer pairs were used to amplify three distinct portions of the 16S gene (the V3, V4, and V4-V5 ranges), making the sequencing data for these samples more complex than for those samples from the other datasets. The Fouhy samples are each a unique combination of either BEI mock community B (16S DNA spike-ins) or BEI mock community C (cultured cells), prepared using one of three library prep protocols, amplified with PCR primers for either the V1-V2 or the V4-V5 region of the 16S gene, and sequenced either on an Illumina MiSeq machine or a Thermo Fisher Ion Torrent. Finally, the 5 Karstens samples originate from a single custom mock DNA library of 8 species, with the V4 region amplified and sequenced on an Illumina MiSeq device.

### 16S amplicon sequencing analysis pipelines

We evaluated four analysis pipelines applied to the 136 mock community samples: Qiime 2, Mothur, PathoScope, and Kraken.

For all Qiime 2 analyses, we used the DADA2 plugin to cluster sequences and construct feature tables. We decided to use the DADA2 plugin rather than its standalone package due to the broad user base of Qiime 2. All mock datasets could be run with paired-end reads besides the Fouhy datasets. In most cases, DADA2 did not require truncation of paired-end sequences, and only the initial 6 bp were trimmed from each read. However, quality scores at the end of nine samples from the Kozich dataset were universally low enough (cutoff of median quality score <20) to require truncation to 240 bp for forward reads and 200 bp for reverse reads for Kozich samples. Taxonomy was assigned using custom naïve Bayes classifiers constructed for each set of mock community samples based on their amplified 16S region. The Qiime artifact output files were converted into BIOM format and subsequently into tab-delimited text format for downstream analyses and pipeline comparisons.

For Mothur analyses, all recommended procedures were followed according to Mothur documentation where possible. For paired-end sequences, the native make.contigs() function was used to join reads. In the pre.cluster() step of Mothur analysis, the “diffs” parameter (the number of mismatches allowed between a cluster’s representative sequence and each member sequence) was set to 2 for joined sequencing reads shorter than 250 bp, 3 for joined reads of length 250-349 bp, and 4 for longer joined reads. For cluster.split(), we set the “taxlevel” parameter to 4, with a “cutoff” of 0.03.

For PathoScope 2.0 analyses, Bowtie2 alignment parameters were set to “--local -R 2 -N 0 -L 25 -i S,1,0.75 -k 10 --score-min L,100,1.28.” These values were optimized for 16S sequencing reads, requiring higher similarity to a reference genome to be considered a hit than the default settings due to the highly conserved nature of portions of the 16S gene. Phylogeny for each taxon was inferred from the NCBI taxon id (ti) for each reference genome using the entrez_fetch() function from the R package *rentrez*.

For Kraken 2 analyses, Kraken taxonomic reports were created for each sample. These were parsed into a taxon/feature counts matrix that included the complete phylogeny for each identified taxon as reported by Kraken 2.

### Bacterial genomic and 16S reference libraries

We used five bacterial sequence reference databases in conjunction with the aforementioned pipelines: Greengenes 13_8, Silva 138, two versions of RefSeq’s representative genomes, and the Kraken Standard library (downloaded on August 20, 2020). According to the Kraken manual, the Kraken Standard library is compiled using the RefSeq database, so it could be considered analogous to the RefSeq2020 library. The RefSeq libraries were downloaded on November 2, 2018, and June 23, 2020; these are denoted as “RefSeq2018” and “RefSeq2020.” Greengenes and Silva are specifically 16S reference databases as they include only sequences for the bacterial 16S gene. RefSeq2018, RefSeq2020, and the Kraken Standard database are all whole-genome libraries, with no special modifications for use with 16S amplicon sequencing data.

### Analysis pipeline and reference library pairings

We analyzed 136 mock community samples using a total of eight distinct pairings of analysis tools and reference libraries: Qiime 2 only with Greengenes (its default reference library), Mothur only with Silva (the default reference library), PathoScope using Greengenes, Silva, RefSeq2018, and RefSeq2020, and Kraken with both its Standard library and with Greengenes. While the Silva database includes species-level taxonomic information for most of its representative 16S sequences, note that Mothur collapses feature counts into genus-level clades and thus does not make species-level calls. The adaptations of Silva used for Kraken 2 and Qiime 2 did not provide species-level calls.

### Tracking available taxonomic information for each ASV/OTU

A counts matrix was created from the results of each of the eight pipeline/reference pairs for each operational taxonomic unit (OTU), amplicon sequence variant (ASV), and feature. Each feature was assigned phylum, class, order, family, genus, species, and subspecies level information when available. Whenever a taxonomic label was missing, the lowest level taxonomy available for a feature was propagated, taking note of what granularity was available (taxonomic best hit). For example, a feature assigned only as a member of the *Bacillales* order would be given the metadata: “phylum: *Firmicutes*, class: *Bacilli*, order: *Bacillales*, family: o_*Bacillales*, genus: o_*Bacillales*, species: o_*Bacillales*.”

### Assessing taxon sensitivity, read specificity, error, and diversity

Several metrics were used in assessing the overall quality and power of each 16S analysis pipeline and reference library at each taxonomic level. Results were independently evaluated at each taxonomic level. Any reads or features not assigned to a taxon at a given phylogenetic level were excluded from analysis, except where otherwise specified.

#### Taxon detection sensitivity

The metric of taxon detection sensitivity is defined here as the portion of expected taxa in a mock community sample detected by a given pipeline, at a minimum of 0.1% relative abundance. It essentially examines how often a given method can correctly determine an organism’s presence in the mock community.

#### Read assignment specificity

Read assignment specificity is defined here as the portion of reads from a given sample assigned to taxa that are actually present in that sample’s mock community. It identifies the frequency of read assignment to incorrect organisms for a given method.

#### Error rate

The Normalized Root Mean Squared Error (NRMSE) was calculated as the root mean square normalized with the assumption that the variance could be increasing given higher read counts. For each sample’s results, given by the equation

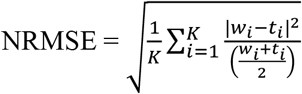

where, for *K* taxa, *w*_*i*_ and *t*_*i*_ are the measured and true read counts of taxon *i*.

#### Alpha diversity estimations

To assess each pipeline’s ability to estimate the true alpha diversity within a sample regardless of accurate species identification, we calculated the log-fold change between the expected and the measured alpha diversity, as measured by the Shannon index, the Simpson index, and the Chao1 index. The R packages *vegan*^*35*^ and *fossil*^*36*^ were used to calculate the Shannon and Simpson indexes, and the Chao1 index, respectively. Due to the alpha diversity metrics’ sensitivity to library and count size differences, we converted the relative abundances of the mock community samples’ ground truths to virtual sequencing libraries of 1,000,000 reads. A rarefaction depth of 10,000 reads per sample was used to normalize all samples and ground truth libraries.

### Statistical methods for significance testing

A series of linear mixed-effects models (LMMs), coupled with *post hoc* least-square means tests and a Tukey multiple comparison correction, were used to determine which pipelines outperformed each other in sensitivity, specificity, error rates, and alpha diversity estimates. LMMs were estimated using the lmer() function from the R package lme4^37^, and *post hoc* comparisons were performed with the lsmeans() function from the R package lsmeans package^38^. These LMMs examine the relevant performance metric as the measured variable, using the 136 mock community samples as a random effect and the pipeline/reference library pair as a fixed effect.

## Results

### Visual evaluation of species-level detection and abundance estimates

**Figure 1** shows stacked bar charts of the results from the Kozich dataset for the ground truth versus all methods at the species level. Overall, pipelines using the Greengenes database (Kraken, Qiime, and Pathoscope) performed the worst in classifying species. PathoScope made the best use of the Greengenes database with the fewest misclassified reads and most correct species-level detection. Kraken (paired with its Standard library) and PathoScope (paired with the RefSeq and Silva libraries) performed best on these datasets. A more quantitative evaluation of these methods in the context of all samples follows.

**Figure 1.**
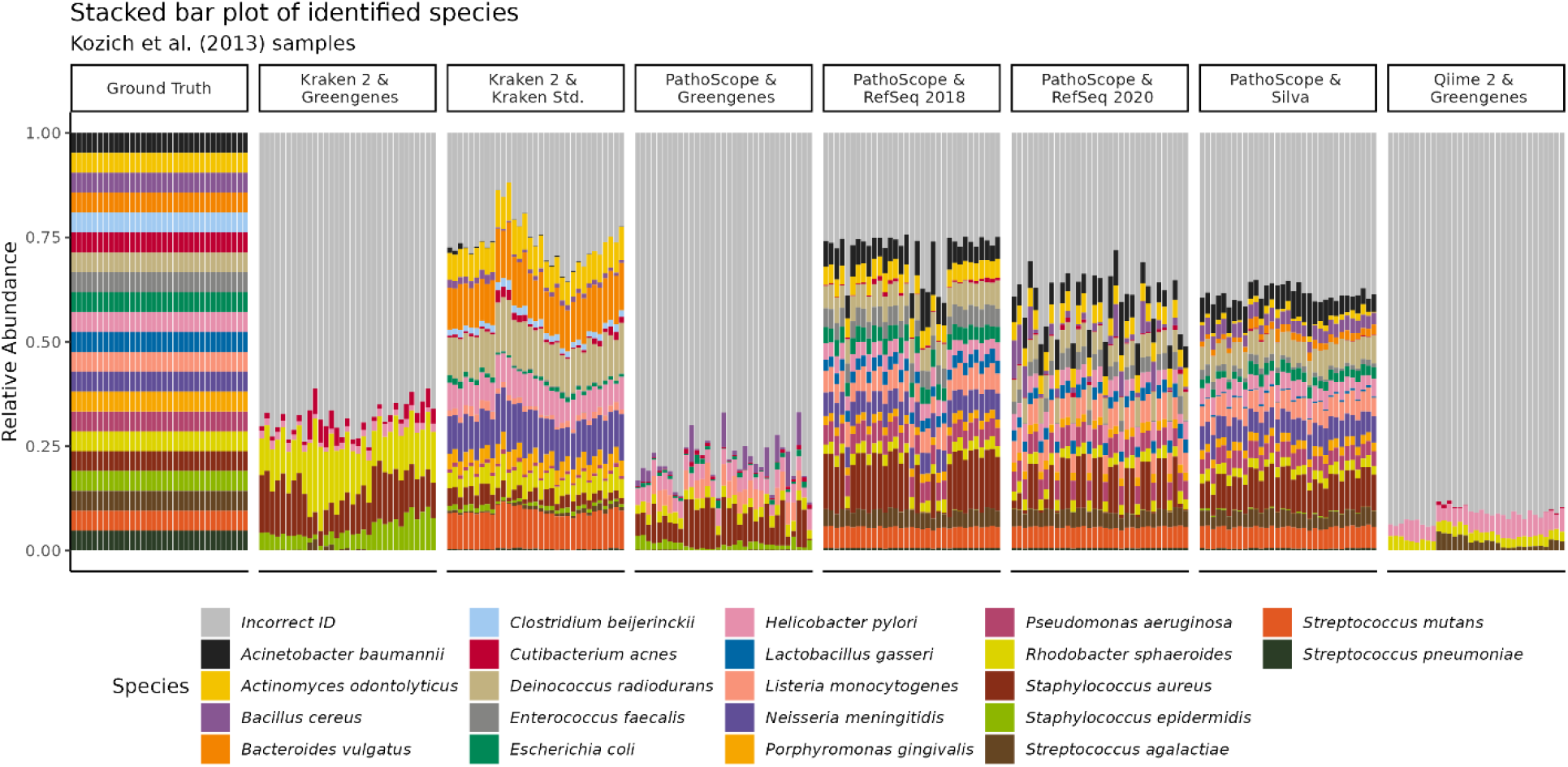
Expected vs measured relative abundances of mock bacteria. A stacked bar plot of the measured relative abundances of bacterial species in 33 samples from Kozich et al. (2013). These samples were all equimolar concentrations of 16S rDNA from 21 species, as shown in the ‘Ground Truth’ bar on the left. All reads assigned to bacterial species other than the 21 expected in the mock community are colored gray and are labeled “Incorrect ID”. Mothur calls were not included as the pipeline does not make species-level calls.

### Taxon detection sensitivity

At the genus level (**Figure 2A**), methods which utilized the Greengenes library were collectively the least sensitive (Qiime 2: mean=0.73, SD=0.16; Kraken: mean=0.73, SD=0.17; PathoScope: mean=0.77, SD=0.24; see Supplemental Table 2 for p-values). When paired with the Silva, RefSeq2018, or RefSeq2020 reference libraries, PathoScope was more sensitive at detecting genera than any other method, peaking when paired with the RefSeq 2018 reference library (mean=0.88, SD=0.14).

**Figure 2.**
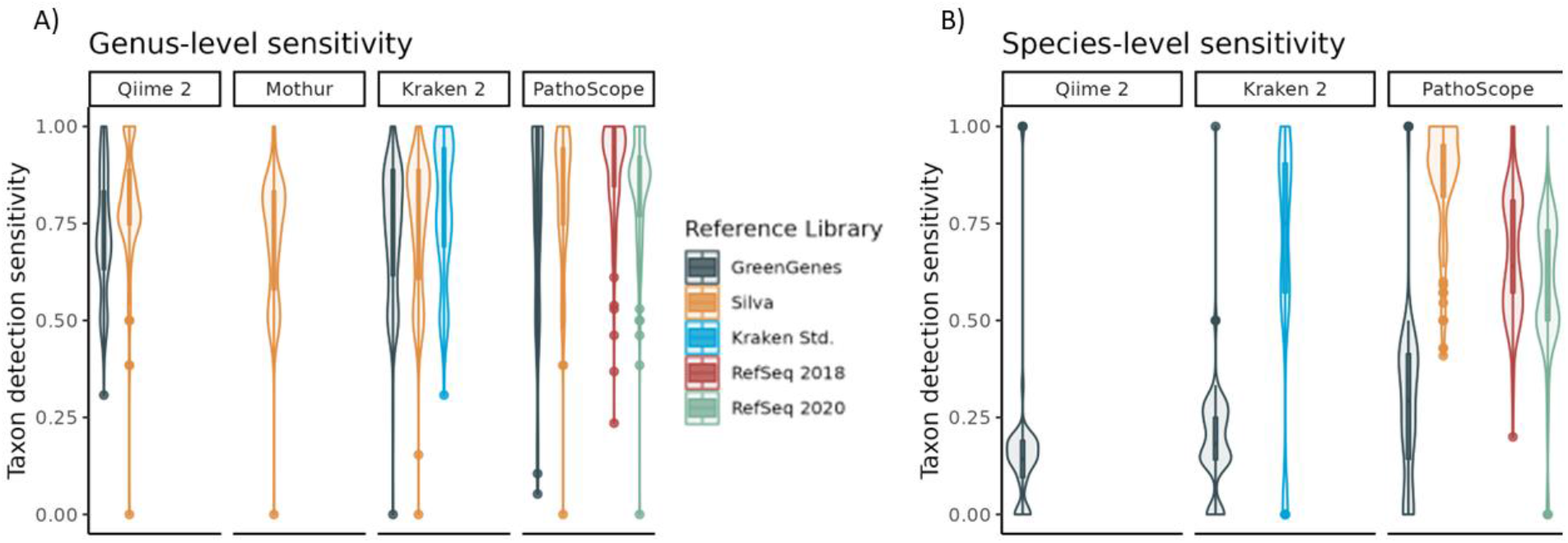
Taxon detection sensitivity of 16S analysis pipelines. Violin plots of the sensitivity of each analysis pipeline and reference library pair used to analyze 16S samples, calculated at the A) genus, and B) species levels. Sensitivity is calculated as the portion of expected taxa in each mock community sample that was detected with least 0.1% relative abundance.

Generally, taxon detection sensitivity was lower at the species level than at the genus level (**Figure 2B**). Methods using Greengenes had extremely low species-level sensitivities (Qiime: mean=0.16, SD=0.18; Kraken: mean=0.18, SD=0.13; PathoScope: mean=0.28, SD=0.21), significantly lower than all other methods (see Supplemental Table 3 for pairwise p-values). Among those methods that used Greengenes, PathoScope was significantly more sensitive than either Qiime or Kraken. The most sensitive method at the species level was PathoScope using the Silva reference library (mean=0.86, SD=0.15), followed by PathoScope using RefSeq2018 (mean=0.67, SD=0.16). Only three species were not detected by PathoScope at a minimum of 0.1% relative abundance in any samples when using Silva as a reference library; these were *Bifidobacterium adolescentis, Prosthecobacter fusiformis*, and *Clostridium beijerinckii*.

### Read assignment specificity

At the genus level, the average read assignment specificity was generally lower for Qiime with GreenGenes, PathoScope with GreenGenes, and Kraken with Silva and its Standard library (**Figure 3A**). However, no overall trends arose in pairwise tests among pipelines and database pairings (see Supplemental Table 4 for pairwise p-values). PathoScope with the RefSeq2018 library (mean=0.91, SD=0.15) and Kraken with GreenGenes (mean=0.89, SD=0.18) had the overall highest read assignment specificity.

**Figure 3.**
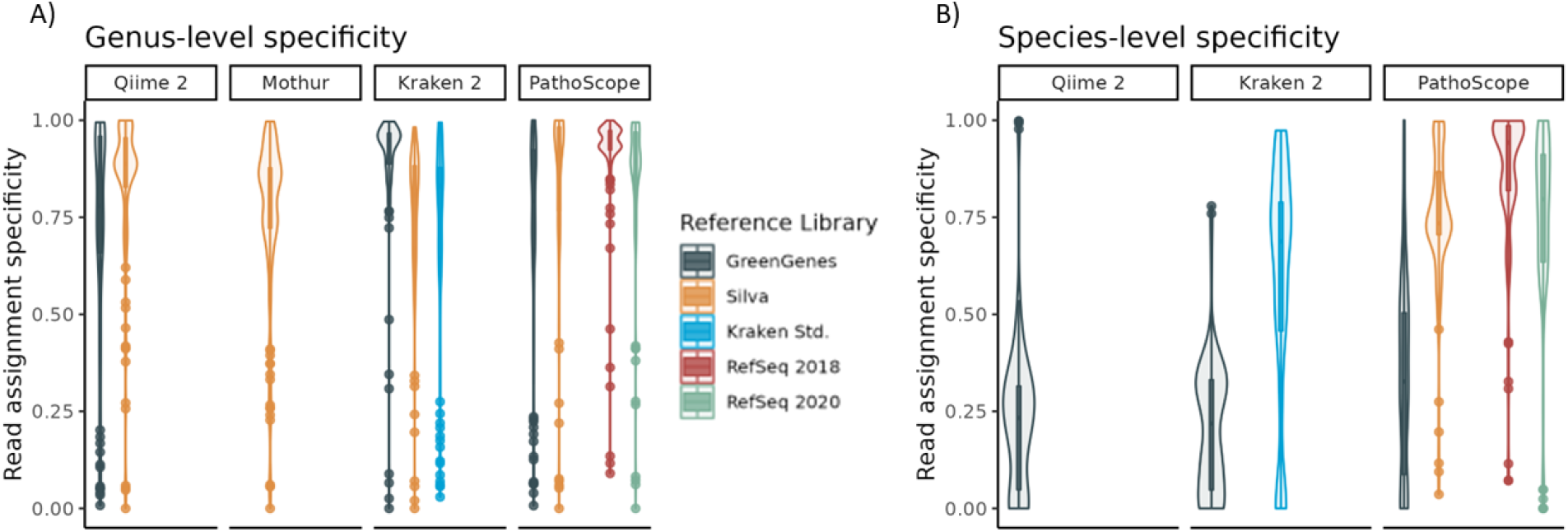
Read assignment specificity of 16S analysis pipelines. Violin plots of the specificity of each analysis pipeline and reference library pair used to analyze 16S samples, calculated at the A) genus and B) species levels. Specificity is calculated as the portion of reads assigned to taxa that are expected to exist within each mock community.

At the species level, both Kraken and Qiime paired with Greengenes had the lowest read assignment specificity (Kraken: mean=0.21, SD=0.17; Qiime 2: mean=0.23, SD=0.2), which were significantly lower than all methods (see Supplemental Table 5 for pairwise p-values). PathoScope, when paired with either the Silva library (mean=0.75, SD=0.18), RefSeq2020 (mean=0.75, SD=0.24), or RefSeq2018 (mean=0.86, SD=0.18) was significantly more specific than Qiime and Kraken (**Figure 3B)**.

### Normalized root mean-square error

Kraken has the lowest error rates, measured as the NRMSE of the raw reads, of all methods evaluated at the genus level regardless of the reference library used (Greengenes: mean=74.02, SD=46.69; Standard: mean=62.95, SD=36.13, Silva: mean=50.41, SD=29.7; see Supplemental Table 6 for pairwise comparison p-values). These were significantly lower than all other error rates. Qiime had the highest genus-level NRMSE for both the Silva and Greengenes libraries (Silva: mean=242.6, SD=115.82; Greengenes: mean=240.78, SD=114.16) of all methods (**Figure 4A**).

**Figure 4.**
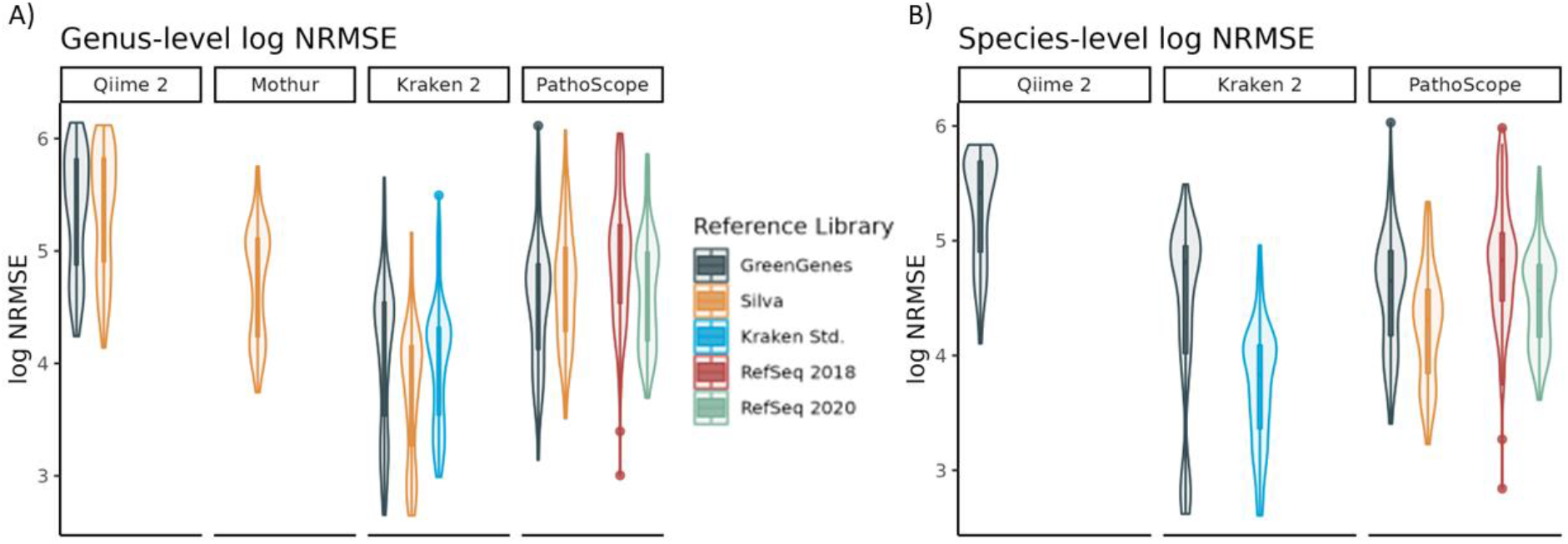
NRMSE of 16S analysis pipelines. Violin plots of the log NRMSE of each analysis pipeline and reference library pair used to analyze 16S samples, calculated at the A) genus, and B) species levels.

Kraken also had the lowest NRMSE at the species level for its Standard database, which was better than all other methods (Standard: mean=47.94; SD=23.88). This was followed by PathoScope for the Silva and RefSeq2020 databases (Silva: mean=79.65, SD=39.35; RefSeq2020: mean=99.83, SD=44.96), and Kraken using Greengenes (mean=100.331, SD=57.23). The worst NRMSE was again held by Qiime using the GreenGenes database (mean=218.3, SD=85.41), which was significantly better than all other methods.

### Alpha diversity estimations

Out of all eight methods evaluated, Kraken paired with Greengenes showed the greatest deviations from expected Shannon (deviation mean=1.05, SD=1.06) and Simpson (deviation mean=0.25, SD=0.27) alpha diversity indexes, significantly higher deviations than all other methods (Tukey-adjusted p<0.001 in all pairwise comparisons). PathoScope generally matched the true Shannon index more closely than all other methods when paired with either RefSeq2018 or RefSeq2020 (RefSeq2018: deviation mean=0.27, SD=0.28; RefSeq2020: deviation mean=0.21, SD=0.23; **Figure 5A**).

**Figure 5.**
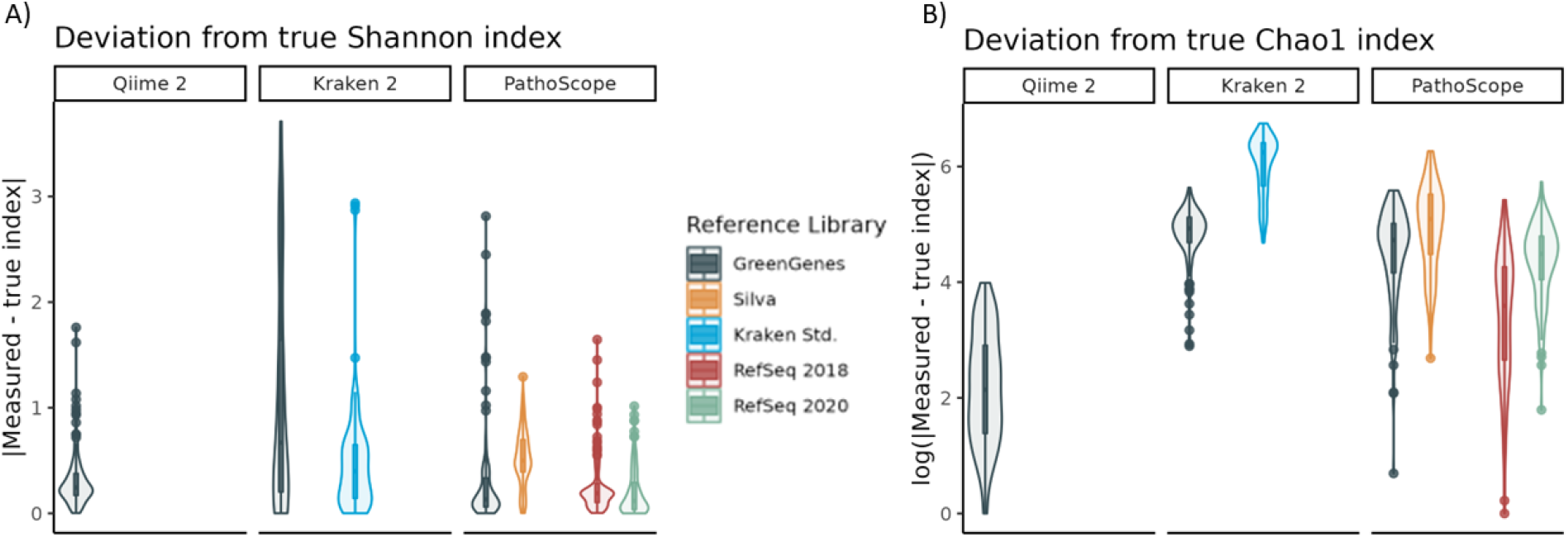
Deviation from true alpha diversity metrics. The absolute difference between the measured Shannon alpha diversity index and the Shannon index value for the true mock community composition, and B) the log of one plus the absolute difference in Chao1 richness estimates and the true number of species present in each mock community. In both cases, values closer to 0 indicate more accurate estimation of the alpha diversity within a sample.

The Chao1 index estimates a sample’s species richness. It roughly amounts to a count of the number of species one would expect to find at a given sequencing depth, which in this case is 10,000 reads. Qiime reported the most closely matching Chao1 indexes, averaging significantly less deviation from the true number of species present (Silva: mean=11.46, SD=14.82; Greengenes: mean=12.47, SD=12.26) than other methods (Tukey-adjusted p<0.001 in all pairwise comparisons). On the other hand, Kraken using its Standard library and Silva frequently overestimated the number of species present by several orders of magnitude (Standard: mean=460.46, SD=191.18; Silva: mean=344.79, SD=135.59), performing worse than all other methods (**Figure 5B**).

Overall, no single pipeline or reference library performed the best in all evaluative metrics, but some holistic trends are present, especially at the species level. **Figure 6B** shows that sensitivity and specificity are correlated traits at the species level (Pearson’s *r* = 0.86) and that PathoScope (regardless of reference library) and Kraken (with its Standard library) dominate the upper right quadrant, where sensitivity and specificity are both high. In particular, PathoScope excels in both sensitivity and specificity when used with either Silva or RefSeq2018. Similarly, **Figure 6C** shows that error and alpha diversity estimation deviation are inversely correlated (Pearson’s r = -0.60) and that no single method yields the lowest alpha diversity deviation and error rates.

**Figure 6.**
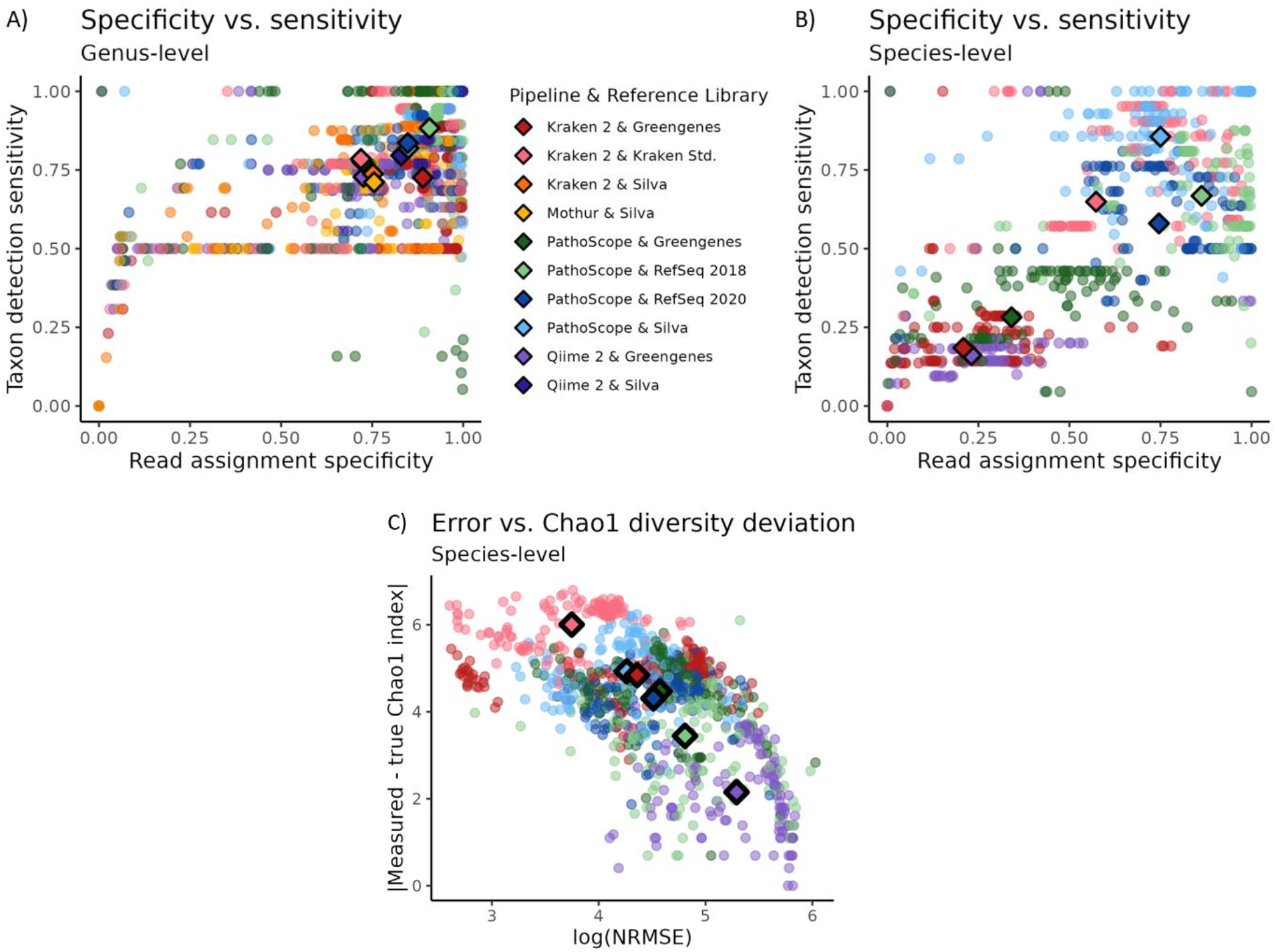
Combined quality of 16S analysis methods. Scatterplots of relative metrics for each 16S analysis pipeline, at the genus (A) and species (B, C) levels. Each point represents a single method’s results when analyzing a single mock community sample. Points are colored by the analysis pipeline/reference library used. Centroids representing the mean values for each pipeline/reference library pair are marked with bolded diamonds.

## Discussion

Mock bacterial communities, either derived from spike-in DNA sequences or extracted from mixtures of bacterial cell monocultures, provide a semblance of a “ground truth” to assess 16S amplicon sequencing analysis methods. Ideally, knowing which species at what amounts should be present in any genuine microbiome sample would allow for accurate identification in every analysis. There are, of course, complications in sequencing experiments: technical bias and errors are introduced into samples at every step of the experiment until being safely sealed as bits in a FASTQ file on a server. Mock species’ relative abundances may be affected by subtle variations in pipetting technique as spike-in DNA is aliquoted from individual sources. Spike-in DNA might be cloned from mutated DNA, or an early PCR error may have propagated through an entire commercial stock of nucleic acids. Different species of bacteria vary in lysing difficulty^39^, causing some species to be underrepresented or even absent in the collected cDNA libraries from a plate^40^. While 16S amplification primers are designed to bind to universal conserved regions of the 16S gene, there is still clearly some amplification bias during PCR^41^. Contamination from reagents^42^, local bacteria in the air, on gloves, or in a pipette tip box can further complicate matters. Thus, limitations of different experimental conditions and methods can dramatically affect the quality of results obtained from mock communities. Present sequencing errors and contamination suggest that amplicon reads will not identify with taxa as precisely as might a tidy, evenly distributed set of sequences drawn from a closed set of well-characterized species. It should then be evident that no analysis pipeline could theoretically exist to perfectly measure a mock community. Such a feat would require identifying only the expected species in their exact proportions, with no extraneous observations. As such, the proximate best method to analyze 16S amplicon sequencing data is one that identifies the makeup of the microbiome as truthfully as possible. Mock microbial communities can provide a level testing ground for existing tools to find their relative strengths and weaknesses in performance.

Of the pipelines tested, both Qiime 2 and Mothur were designed and built specifically for 16S amplicon sequencing analysis. Each has a suite of utility functions built to assist researchers in processing their data from the sequencer to differential abundance analysis and visualizations. Both are typically bundled with a dedicated bacterial 16S gene sequence database’s reference library for alignment (i.e., Greengenes for Qiime 2, Silva for Mothur). However, our results present strong evidence that PathoScope and Kraken 2 outperform both tools, even when comparing reads to identical reference databases. This phenomenon interestingly occurs despite Kraken 2 and PathoScope’s status as more general whole genome sequencing and metagenomics data tools. In pairwise comparisons, PathoScope is more sensitive and specific at taxon detection and has a lower error score than either Qiime 2 (when both tools use Greengenes) or Mothur (when both tools use Silva) and has comparable alpha diversity index estimates at both the genus and species levels. In general, Silva’s outperformance of Greengenes confirmed results found in previous benchmarks of 16S amplicon sequencing analysis methods^19-22,43^.

Kraken 2, when used with its Standard library, was rarely the top-performing analytic method in terms of sensitivity or specificity, although it was generally less error-prone than Qiime, Mothur, or any tool using Greengenes as a reference library. Kraken 2 has the added practical utility of being extremely fast and easy to use. Yet one constraint in analyzing Kraken results is that they cannot be upsampled from a given taxon level, whereas PathoScope, Qiime, and Mothur all allow the tracing back of the taxonomic hierarchy of a given microbe. Both Qiime 2 and Mothur take advantage of naïve Bayes classifiers, which work most efficiently when trained on the specific region of the 16S gene amplified by PCR primers.Overall, PathoScope was the most sensitive in detecting taxa and specific in assigning reads, and the least error-prone tool when paired with either Silva or RefSeq2018. However, it was not without limitations, as its computational expenses seemed to be an additional order of magnitude above those of other methods. This was evident from large interim SAM files (>128GB) and runtimes on the order of several hours, whereas Kraken 2 in particular took mere minutes. Issues aside, PathoScope is likely to outperform Qiime 2 and Mothur in identification regardless of the database used. This finding partly results from PathoScope’s Bayesian mixed modeling identification algorithm, which accounts for the possibility that multiple species can be present in the sample or that the target strain is not present in the reference database. PathoScope consistently outperformed Kraken in most cases, although the difference was often slight and not statistically significantly better. Overall, these comparisons show that methods designed for general metagenomics analyses consistently outperform methods specifically designed for analyzing 16S data.

While many species are identifiable from their 16S gene sequence or a single hypervariable region, it is important to note that imperfect accuracy at this level is not solely a computational issue. For example, although the 16S rRNA gene is approximately 1550 bp long, the short sequencing reads obtained in most next-generation sequencing (NGS) only span about 250–500 bases and lack ideal resolution at the species level^44^. Compared to NGS, long-read sequencing technologies have been shown to perform better in classifying at the genus and species level^45,46^. Furthermore, a major limitation to 16S amplicon studies is that some clades of bacteria exist with identical 16S DNA in the commonly sequenced V4 region. These clades of difficult-to-identify bacteria make up the bulk of Kraken’s and PathoScope’s incorrect calls. For example, *Bifidobacterium adolescentis* was nearly universally misclassified by all methods as other *Bifidobacterium* species, and *Prosthecobacter fusiformis* was frequently misidentified as *Prosthecobacter dejongeii*, a species with which it shares over 99% of its 16S DNA sequence^47^. Even further complications arise from many bacteria having several copies of the 16S gene, which may not be identical between operons within a genome^48^. This latter point may be in part why metagenomic methods such as Kraken 2 and PathoScope outperform specific methods such as Qiime 2 and Mothur, especially at the species level. The metagenomic methods are better designed to take into account multiple 16S genes if present.

One of PathoScope’s largest sources of error and lost taxon detection sensitivity calls when using the RefSeq2020 library comes from an apparent erroneous reference genome scaffold in the RefSeq representative genomes. In all mock community samples containing *Escherichia coli*, PathoScope with RefSeq2020 reported the presence of *Tumebacillus flagellates* at relative abundances tightly correlated with the expected values of *E. coli* (Pearson’s *r* = 0.959). The circumstances strongly imply that reads actually originating from *E. coli* were incorrectly assigned to *T. flagellates. T. flagellates* is not even in the same phylum as *E. coli*, so the casual misassignment of reads between the species would be extremely unlikely based on 16S sequence similarity. Instead, a pairwise BLAST comparing *E. coli*’s 16S gene sequence to the *T. flagellates* scaffolds using the exact RefSeq entry that PathoScope had assigned those reads to (accession: NZ_JMIR01000093)^49^ revealed that one *T. flagellates* scaffold had a 100% identity alignment over 911 bp. The finding possibly represents a case of horizontal gene transfer of the 16S gene, but it appears far more likely that *E. coli* contamination existed in the DNA library, which then was sequenced and assembled into *T. flagellates* scaffolds. On further study, it became apparent that this is merely one example of pervasive sequence contamination, meaning the accidental inclusion of sequences from other organisms or the misclassification of sequences, in public genome databases. This phenomenon has been recently explored in the NCBI RefSeq database^50-52^. The recent prevalence of high throughput and the accelerating low cost of next-generation sequencing (NGS) technologies has led to a rapid increase in published genomes available in the RefSeq libraries, although imperfect methods and protocols for sequencing data are contributing to high contamination rates. Human contamination in published genomes, while not a problem in 16S analyses, is a particularly frustrating problem when analyzing shotgun metagenomics data. Clearly, metagenomic read-mapping approaches such as Kraken and PathoScope afford the potential for the development of novel quality control pipelines for RefSeq and other genome sequence databases.

The increasing prevalence of poor sequencing quality control helps to explain why the RefSeq 2018 libraries often performed better than the 2020 libraries. Many tools have been developed to identify and correct contamination errors in sequences and public databases^52-56^, but this is an ongoing problem that demands additional filtering and correction efforts after directly retrieving libraries from the public repository. Given the higher specificity and sensitivity of PathoScope when using the 2018 RefSeq library over the 2020 library, we recommend using older RefSeq libraries until newer versions have been processed to remove contamination. Also of interest to note is the high accuracy of Silva in its species calls when using PathoScope, even though it cannot be used to make such calls when used with Qiime 2, Mothur, or Kraken 2. Silva also presents as a viable alternative to the RefSeq libraries in avoiding contamination.

## Conclusion

While both Qiime 2 and Mothur excel at assigning family-level or higher taxonomy to 16S amplicon sequences, they struggle to maintain accuracy at the genus level or more granular taxonomic analyses. Kraken 2, despite its primary purpose for whole genome sequencing metagenomics analyses, offers more power in analyzing 16S data without any increase in computational costs. PathoScope, while computationally more expensive, produces the most sensitive and accurate results of all evaluated pipelines when used on a diverse set of mock bacterial community samples. Analysis pipelines using Silva as a reference library outperformed those using Greengenes, and PathoScope using Silva yielded the highest accuracies and sensitivities. While whole-genome reference libraries, such as Kraken’s Standard or RefSeq’s representative genomes, may provide some benefits over Silva in terms of sensitivity, they may yield more spurious species-level calls. Based on the research conducted here with mock microbial communities, we recommend Silva and RefSeq above other databases and discourage usage of the Greengenes reference library for future analysis. We also recommend PathoScope and Kraken as fully capable, competitive options for conducting genus- and species-level 16S amplicon sequencing data analysis, in addition to outperforming other tools when using shotgun metagenomics data^17^.

## Availability of data and materials

Reference libraries used in analysis are available in the following GitHub repository: https://github.com/aubreyodom/16SBenchmarking.

## Bibliography

1. Johnson JS, Spakowicz DJ, Hong BY, et al. Evaluation of 16S rRNA gene sequencing for species and strain-level microbiome analysis. Nat Commun. 11 06 2019;10(1):5029. doi:10.1038/s41467-019-13036-1

2. Callahan BJ, McMurdie PJ, Holmes SP. Exact sequence variants should replace operational taxonomic units in marker-gene data analysis. ISME J. 12 2017;11(12):2639–2643. doi:10.1038/ismej.2017.119

3. Callahan BJ, McMurdie PJ, Rosen MJ, Han AW, Johnson AJ, Holmes SP. DADA2: High-resolution sample inference from Illumina amplicon data. Nat Methods. 07 2016;13(7):581–3. doi:10.1038/nmeth.3869

4. Bolyen E, Rideout JR, Dillon MR, et al. Reproducible, interactive, scalable and extensible microbiome data science using QIIME 2. Nat Biotechnol. 08 2019;37(8):852–857. doi:10.1038/s41587-019-0209-9

5. Caporaso JG, Kuczynski J, Stombaugh J, et al. QIIME allows analysis of high-throughput community sequencing data. Nat Methods. May 2010;7(5):335–6. doi:10.1038/nmeth.f.303

6. Schloss PD, Westcott SL, Ryabin T, et al. Introducing mothur: open-source, platform-independent, community-supported software for describing and comparing microbial communities. Appl Environ Microbiol. Dec 2009;75(23):7537–41. doi:10.1128/AEM.01541-09

7. Kopylova E, Navas-Molina JA, Mercier C, et al. Open-Source Sequence Clustering Methods Improve the State Of the Art. mSystems. 2016 Jan-Feb 2016;1(1) doi:10.1128/mSystems.00003-15

8. Westcott SL, Schloss PD. De novo clustering methods outperform reference-based methods for assigning 16S rRNA gene sequences to operational taxonomic units. PeerJ. 2015;3:e1487. doi:10.7717/peerj.1487

9. Edgar RC. Updating the 97% identity threshold for 16S ribosomal RNA OTUs. Bioinformatics. 07 15 2018;34(14):2371–2375. doi:10.1093/bioinformatics/bty113

10. Amir A, McDonald D, Navas-Molina JA, et al. Deblur Rapidly Resolves Single-Nucleotide Community Sequence Patterns. mSystems. 2017 Mar-Apr 2017;2(2) doi:10.1128/mSystems.00191-16

11. Hong C, Manimaran S, Shen Y, et al. PathoScope 2.0: a complete computational framework for strain identification in environmental or clinical sequencing samples. Microbiome. 2014;2:33. doi:10.1186/2049-2618-2-33

12. Francis OE, Bendall M, Manimaran S, et al. Pathoscope: species identification and strain attribution with unassembled sequencing data. Genome research. 2013;23(10):1721–1729.

13. Byrd AL, Perez-Rogers JF, Manimaran S, et al. Clinical PathoScope: rapid alignment and filtration for accurate pathogen identification in clinical samples using unassembled sequencing data. BMC bioinformatics. 2014;15(1):1–14.

14. Wood DE, Lu J, Langmead B. Improved metagenomic analysis with Kraken 2. Genome Biol. 11 28 2019;20(1):257. doi:10.1186/s13059-019-1891-0

15. He Y, Caporaso JG, Jiang XT, et al. Stability of operational taxonomic units: an important but neglected property for analyzing microbial diversity. Microbiome. 2015;3:20. doi:10.1186/s40168-015-0081-x

16. Nearing JT, Douglas GM, Comeau AM, Langille MGI. Denoising the Denoisers: an independent evaluation of microbiome sequence error-correction approaches. PeerJ. 2018;6:e5364. doi:10.7717/peerj.5364

17. Miossec MJ, Valenzuela SL, Pérez-Losada M, Johnson WE, Crandall KA, Castro-Nallar E. Evaluation of computational methods for human microbiome analysis using simulated data. PeerJ. 2020;8:e9688.

18. Miossec MJ, Valenzuela SL, Mendez KN, Castro-Nallar E. Computational methods for human microbiome analysis. Current Protocols in Microbiology. 2017;47(1):1E. 14.1-1E. 14.17.

19. Dixit K, Davray D, Chaudhari D, et al. Benchmarking of 16S rRNA gene databases using known strain sequences. Bioinformation. 2021;17(3):377–391. doi:10.6026/97320630017377

20. López-García A, Pineda-Quiroga C, Atxaerandio R, et al. Comparison of Mothur and QIIME for the Analysis of Rumen Microbiota Composition Based on 16S rRNA Amplicon Sequences. Front Microbiol. 2018;9:3010. doi:10.3389/fmicb.2018.03010

21. Almeida A, Mitchell AL, Tarkowska A, Finn RD. Benchmarking taxonomic assignments based on 16S rRNA gene profiling of the microbiota from commonly sampled environments. Gigascience. 05 01 2018;7(5) doi:10.1093/gigascience/giy054

22. Lu J, Salzberg SL. Ultrafast and accurate 16S rRNA microbial community analysis using Kraken 2. Microbiome. 08 28 2020;8(1):124. doi:10.1186/s40168-020-00900-2

23. DeSantis TZ, Hugenholtz P, Larsen N, et al. Greengenes, a chimera-checked 16S rRNA gene database and workbench compatible with ARB. Appl Environ Microbiol. Jul 2006;72(7):5069–72. doi:10.1128/AEM.03006-05

24. Quast C, Pruesse E, Yilmaz P, et al. The SILVA ribosomal RNA gene database project: improved data processing and web-based tools. Nucleic Acids Res. Jan 2013;41(Database issue):D590–6. doi:10.1093/nar/gks1219

25. Cole JR, Wang Q, Fish JA, et al. Ribosomal Database Project: data and tools for high throughput rRNA analysis. Nucleic Acids Res. Jan 2014;42(Database issue):D633–42. doi:10.1093/nar/gkt1244

26. Lappan R, Imbrogno K, Sikazwe C, et al. A microbiome case-control study of recurrent acute otitis media identified potentially protective bacterial genera. BMC Microbiol. 02 20 2018;18(1):13. doi:10.1186/s12866-018-1154-3

27. De Boeck I, Wittouck S, Wuyts S, et al. Comparing the Healthy Nose and Nasopharynx Microbiota Reveals Continuity As Well As Niche-Specificity. Front Microbiol. 2017;8:2372. doi:10.3389/fmicb.2017.02372

28. Lapidot R, Faits T, Ismail A, et al. Nasopharyngeal Dysbiosis Precedes the Development of Lower Respiratory Tract Infections in Young Infants: a longitudinal infant cohort study. medRxiv. 2021;

29. Schoch CL, Ciufo S, Domrachev M, et al. NCBI Taxonomy: a comprehensive update on curation, resources and tools. Database (Oxford). 01 01 2020;2020 doi:10.1093/database/baaa062

30. O’Leary NA, Wright MW, Brister JR, et al. Reference sequence (RefSeq) database at NCBI: current status, taxonomic expansion, and functional annotation. Nucleic Acids Res. Jan 04 2016;44(D1):D733–45. doi:10.1093/nar/gkv1189

31. Lluch J, Servant F, Païssé S, et al. The Characterization of Novel Tissue Microbiota Using an Optimized 16S Metagenomic Sequencing Pipeline. PLoS One. 2015;10(11):e0142334. doi:10.1371/journal.pone.0142334

32. Kozich JJ, Westcott SL, Baxter NT, Highlander SK, Schloss PD. Development of a dual-index sequencing strategy and curation pipeline for analyzing amplicon sequence data on the MiSeq Illumina sequencing platform. Appl Environ Microbiol. Sep 2013;79(17):5112–20. doi:10.1128/AEM.01043-13

33. Fouhy F, Clooney AG, Stanton C, Claesson MJ, Cotter PD. 16S rRNA gene sequencing of mock microbial populations-impact of DNA extraction method, primer choice and sequencing platform. BMC Microbiol. 06 24 2016;16(1):123. doi:10.1186/s12866-016-0738-z

34. Karstens L, Asquith M, Davin S, et al. Controlling for Contaminants in Low-Biomass 16S rRNA Gene Sequencing Experiments. mSystems. Jun 04 2019;4(4) doi:10.1128/mSystems.00290-19

35. Oksanen J, Kindt R, Legendre P, et al. The vegan package: community ecology package, version 1.13-1 URL: http://vegan r-forge r-project org. 2008;

36. Vavrek MJ. Fossil: palaeoecological and palaeogeographical analysis tools. Palaeontologia electronica. 2011;14(1):16.

37. Bates D, Maechler M, Bolker B, Walker S. Fitting linear mixed-effects models using lme4. Journal of Statistical Software 67: 1–48. 2015.

38. Lenth RV. Least-squares means: the R package lsmeans. Journal of statistical software. 2016;69:1–33.

39. Gill C, van de Wijgert JH, Blow F, Darby AC. Evaluation of Lysis Methods for the Extraction of Bacterial DNA for Analysis of the Vaginal Microbiota. PLoS One. 2016;11(9):e0163148. doi:10.1371/journal.pone.0163148

40. Boers SA, Jansen R, Hays JP. Understanding and overcoming the pitfalls and biases of next-generation sequencing (NGS) methods for use in the routine clinical microbiological diagnostic laboratory. Eur J Clin Microbiol Infect Dis. Jun 2019;38(6):1059–1070. doi:10.1007/s10096-019-03520-3

41. Sze MA, Schloss PD. The Impact of DNA Polymerase and Number of Rounds of Amplification in PCR on 16S rRNA Gene Sequence Data. mSphere. 05 22 2019;4(3) doi:10.1128/mSphere.00163-19

42. Salter SJ, Cox MJ, Turek EM, et al. Reagent and laboratory contamination can critically impact sequence-based microbiome analyses. BMC biology. 2014;12(1):1–12.

43. Straub D, Blackwell N, Langarica-Fuentes A, Peltzer A, Nahnsen S, Kleindienst S. Interpretations of Environmental Microbial Community Studies Are Biased by the Selected 16S rRNA (Gene) Amplicon Sequencing Pipeline. Front Microbiol. 2020;11:550420. doi:10.3389/fmicb.2020.550420

44. Yang B, Wang Y, Qian P-Y. Sensitivity and correlation of hypervariable regions in 16S rRNA genes in phylogenetic analysis. BMC bioinformatics. 2016;17(1):1–8.

45. Nygaard AB, Tunsjø HS, Meisal R, Charnock C. A preliminary study on the potential of Nanopore MinION and Illumina MiSeq 16S rRNA gene sequencing to characterize building-dust microbiomes. Scientific Reports. 2020;10(1):1–10.

46. Pearman WS, Freed NE, Silander OK. Testing the advantages and disadvantages of short-and long-read eukaryotic metagenomics using simulated reads. BMC bioinformatics. 2020;21(1):1–15.

47. Lee J, Park B, Woo SG, Park J. Prosthecobacter algae sp. nov., isolated from activated sludge using algal metabolites. Int J Syst Evol Microbiol. Feb 2014;64(Pt 2):663–667. doi:10.1099/ijs.0.052787-0

48. Louca S, Doebeli M, Parfrey LW. Correcting for 16S rRNA gene copy numbers in microbiome surveys remains an unsolved problem. Microbiome. 02 26 2018;6(1):41. doi:10.1186/s40168-018-0420-9

49. Wang Q, Xie N, Qin Y, et al. Tumebacillus flagellatus sp. nov., an α-amylase/pullulanase-producing bacterium isolated from cassava wastewater. Int J Syst Evol Microbiol. Sep 2013;63(Pt 9):3138–3142. doi:10.1099/ijs.0.045351-0

50. Lupo V, Van Vlierberghe M, Vanderschuren H, Kerff F, Baurain D, Cornet L. Contamination in Reference Sequence Databases: Time for Divide-and-Rule Tactics. Front Microbiol. 2021;12:755101. doi:10.3389/fmicb.2021.755101

51. Breitwieser FP, Pertea M, Zimin AV, Salzberg SL. Human contamination in bacterial genomes has created thousands of spurious proteins. Genome Res. 06 2019;29(6):954–960. doi:10.1101/gr.245373.118

52. Steinegger M, Salzberg SL. Terminating contamination: large-scale search identifies more than 2,000,000 contaminated entries in GenBank. Genome Biol. 05 12 2020;21(1):115. doi:10.1186/s13059-020-02023-1

53. Lu J, Salzberg SL. Removing contaminants from databases of draft genomes. PLoS Comput Biol. 06 2018;14(6):e1006277. doi:10.1371/journal.pcbi.1006277

54. Cornet L, Baurain D. Contamination detection in genomic data: more is not enough. Genome Biol. 02 21 2022;23(1):60. doi:10.1186/s13059-022-02619-9

55. De Simone G, Pasquadibisceglie A, Proietto R, et al. Contaminations in (meta)genome data: An open issue for the scientific community. IUBMB Life. 04 2020;72(4):698–705. doi:10.1002/iub.2216

56. Nasko DJ, Koren S, Phillippy AM, Treangen TJ. RefSeq database growth influences the accuracy of k-mer-based lowest common ancestor species identification. Genome biology. 2018;19(1):1–10.

